# Metabolite-responsive Control of Transcription by Phase Separation-based Synthetic Organelles

**DOI:** 10.1101/2024.09.17.613456

**Authors:** Carolina Jerez-Longres, Wilfried Weber

**Affiliations:** INM – Leibniz Institute for New Materials, Campus D2 2, 66123 Saarbrücken, Germany; Saarland University, Department of Materials Science and Engineering, 66123 Saarbrücken, Germany; Signalling research Centers BIOSS and CIBSS, Faculty of Biology, and SGBM - Spemann Graduate School of Biology and Medicine, University of Freiburg, Schänzlestrasse 18, 79104 Freiburg, Germany

**Keywords:** Coacervate, intrinsically disordered region, in vitro transcription, liquid-liquid phase separation, repressor protein

## Abstract

Living natural materials have remarkable sensing abilities that translate external cues into functional changes of the material. The reconstruction of such sensing materials in bottom-up synthetic biology provides the opportunity to develop synthetic materials with life-like sensing and adaptation ability. Key to such functions are material modules that translate specific input signals into a biomolecular response. Here, we engineer a synthetic organelle based on liquid-liquid phase separation that translates a metabolic signal into the regulation of gene transcription. To this aim, we engineer the pyruvate-dependent repressor PdhR to undergo liquid-liquid phase separation *in vitro* by fusion to intrinsically disordered regions. We demonstrate that the resulting coacervates bind DNA harbouring PdhR-responsive operator sites in a pyruvate dose-dependent and reversible manner. We observed that the activity of transcription units on the DNA was strongly attenuated following recruitment to the coacervates. However, the addition of pyruvate resulted in a reversible and dose-dependent reconstitution of transcriptional activity. The coacervate-based synthetic organelles linking metabolic cues to transcriptional signals represent a materials approach to confer stimulus-responsiveness to minimal bottom-up synthetic biological systems and open opportunities in materials for sensor applications.

## Introduction

Natural living materials show remarkable features such as stimulus-responsiveness or self-organization. Key to stimulus responsiveness of natural materials is the ability of biomolecules to translate external cues such as the concentration of a metabolite into a distinct biochemical output like a change in protein structure, interaction, or enzyme activity. With the emergence of synthetic biology and the ever-growing opportunities of engineering biological systems, it became possible to use such biological receptors to program properties and functions of biological and biohybrid materials^1–3^. For example, the functional coupling of engineered biological receptors to polymers enabled the synthesis of polymer materials that change property and function in response to external cues. Here, the widely occurring mechanism of biological receptors to homo- or heterodimerize in response to an external stimulus has been used to conditionally crosslink (bio-) chemical polymers and thus to change the macroscopic material properties. For example, polymers were crosslinked by receptors that dissociate in response to small molecule drugs^4,5^, which enabled a dose-dependent dissolution of the resulting material and the release of (therapeutic) biomolecular cargo. More recently, engineered photoreceptors previously used in molecular optogenetics haven been used to crosslink polymers such as polyethylene glycol. With this approach, hydrogel materials have been synthesized that change mechanical properties in a reversible and dose-dependent manner in response to multichromatic light^6–10^.

Similar to the modular design of computational synthetic genetic networks by interconnecting genetic switches, individual biological receptors have been integrated into polymer materials and functionally wired via diffusible signals so that the overall material was able to perform fundamental computational operations such as Boolean algebra, signal amplification, or counting^11–13^. While such materials show high promise as smart sensing devices or as extracellular, dynamically tuneable matrix for tissue engineering^6^, programmable biological materials are also key to bottom-up synthetic biology aiming at the assembly of systems with life-like or even living properties. For example, the integration of engineered synthetic biological receptors into bio-based or synthetic particles enabled the design of stimulus-responsive cell-like moieties^14^ or the stimulus-inducible assembly and reorganization of synthetic “tissues” from designer protocells^15–17^.

One important process in biological self-assembly and self-organization is based on liquid-liquid phase separation (LLPS), in which biomolecules spontaneously separate in distinct phases. LLPS is the driving mechanism behind the formation of membrane-less organelles such as nucleoli or Cajal bodies that serve as localized biochemical reaction centres^18^. The primary driving forces for the formation of biomolecular condensates are the concentration of the involved molecules (such as proteins, RNA, or DNA), the biophysical properties of the biopolymers involved, and the multivalent interactions among them. Multivalent interaction is often facilitated by intrinsically disordered regions (IDRs), which are sequences of amino acids lacking a defined secondary structure and that can interact via cation-pi, pi-stacking or electrostatic interactions, for instance^19^. Proteins like Fused in Sarcoma (FUS)^18^ and heterogeneous nuclear ribonucleoprotein A1 (HNRNPA1)^20^ are well-known examples, as they have been documented to undergo phase separation when a critical concentration threshold is reached. LLPS has emerged as a versatile approach for compartmentalization in bottom-up synthetic biology ^21^. For instance, phase separated compartments which exhibit enzymatic activity have been described ^22–27^. In addition, enabled by cell free synthetic biology technologies ^28^, *in vitro* transcription-translation has been achieved in such phase separated condensates ^29–31^.

In this work, we harness the principle of liquid-liquid phase separation to assemble synthetic organelles that are able to integrate metabolic signals and to translate these signals into the control of transcription. The synthetic organelles are based on the pyruvate-responsive repressor PdhR engineered to undergo liquid-liquid phase separation by fusion to an intrinsically disordered region. We demonstrate that the resulting coacervates can sequester DNA harbouring cognate *pdhO* operator sequences in a pyruvate-dependent manner. We further demonstrate that the sequestration of the DNA correlated with strongly inhibited transcription that was reversible in a dose-dependent manner by the addition of increasing pyruvate concentrations.

The reversible and dose-dependent behaviour of these coacervates highlights their potential as dynamic materials for bottom-up synthetic biology, providing a new tool for the regulation of gene expression in response to metabolic signals. Through this work, we aim to advance the understanding of phase-separated systems in bottom-up synthetic biology and pave the way for the development of new materials with applications in biosensing, and beyond.

## Material and methods

### Construction of expression vectors

The sequences of the nucleic acid constructs used in this study are shown in **Table S1**. Bacteria with constructs containing more than one *pdhO* repeat were grown at 30 °C to prevent loss of repeats by recombination.

### Protein production and purification

#### Production of 3C protease

*E. coli* BL21(DE3)-pLysS cells (Thermo Fisher Scientific, Waltham, MA, cat. no. C602003) were transformed with plasmid pHJW257 ^32^, coding for strep-tagged 3CP, and cultivated in Luria/Miller broth (LB) supplemented with ampicillin (100 μg/mL) and chloramphenicol (36 μg/mL). Bacteria were grown in LB medium in flasks at 37 °C while shaking to an OD_600_ of 0.9 prior to induction with 1 mM isopropyl β-D-1-thiogalactopyranoside (IPTG; Carl Roth, Karlsruhe, Germany, cat. no. 2316.5) and a protein production period of 5 h at 37 °C. Subsequently, the bacteria were harvested by centrifugation and the cells were resuspended in column buffer (100 mM Tris-HCl, 150 mM NaCl, pH 8.0, 35 mL per 1 L of initial culture), shock-frozen in liquid nitrogen and stored at -80 °C until purification. For affinity chromatography purification, resuspended pellets were lysed by ultrasonication (Sonoplus HD, Bandelin, Berlin, Germany). The lysates were clarified by centrifugation at 30 000 g for 30 min and the supernatant was loaded onto a gravity-flow column containing StrepTactin XT 4Flow resin (IBA Lifesciences, Göttingen, Germany, cat. no. 2-5030-002; 1.5 mL StrepTactin beads per 1 L bacterial culture) previously equilibrated with column buffer. The column was washed with 10 column volumes (CV) column buffer, after which the protein was eluted in 7 fractions of 1 CV each with elution buffer (100 mM Tris-HCl, 150 mM NaCl, 50 mM biotin, pH 8.0). The fractions with the highest absorbance at 280 nm were pooled and β-mercaptoethanol was added to a final concentration of 10 mM. The eluate was concentrated using a Vivaspin Turbo 5k-molecular weight cut-off (MWCO) spin concentrator (Sartorius AG, Göttingen, Germany, cat. no. VS15T12), and the final concentration was determined by Bradford assay. Finally, 10% (v/v) glycerol was added and the protein was shock-frozen in single-use aliquots in liquid nitrogen and stored at -80 °C.

#### Production of PdhR and PdhR-FUS_N_

*E. coli* BL21(DE3) cells (Thermo Fisher Scientific, cat. no. C601003) were transformed with plasmid pRG001 (PdhR) ^33^ or pCJL236 (MBP-PdhR-FUS_N_), and selected by growth in Luria/Miller broth (LB) supplemented with ampicillin (100 μg/mL). Pre-cultures were grown overnight and were used to inoculate expression cultures in flasks of LB medium supplemented with ampicillin. The latter were incubated at 37 °C while shaking to an OD_600_ of 0.9 prior to induction with 1 mM IPTG and a protein production period of 4 h at 37 °C. The bacteria were harvested by centrifugation and resuspended in lysis buffer (50 mM NaH_2_PO_4_, 300 mM NaCl, 10 mM imidazole, pH 8.0; 35 mL per 1 L of initial culture), shock-frozen in liquid nitrogen and stored at -80 °C until purification. For purification, the pellets were thawed, and 0.5 mM tris(2-carboxyethyl)phosphine (TCEP) was added. The cells were lysed on an APV 2000 French press (APV Manufacturing, Bydgoszcz, Poland) at 1000 bar. The lysate was clarified by centrifugation at 30 000 g for 30 min. For PdhR purification, the clarified lysate was loaded onto a gravity-flow column containing a nickel-nitrilotriaceticacid (Ni-NTA) Superflow agarose resin (Qiagen, Hilden, Germany, cat. no. 30410; 1 mL resin per 1 L culture) previously equilibrated with lysis buffer. The column was washed twice with 10 CV wash buffer (50 mm NaH_2_PO_4_, 300 mm NaCl, 20 mm imidazole, 0.5 mM TCEP, pH 8.0) and the protein was eluted in 8 CV elution buffer (50 mm NaH_2_PO_4_, 300 mm NaCl, 250 mm imidazole, 0.5 mM TCEP, pH 8.0). The MBP-PdhR-FUS_N_ protein was purified using an ÄKTA Explorer fast protein liquid chromatography system (GE Healthcare, Freiburg, Germany). The clarified lysate was loaded on a Ni-NTA agarose column and, after washing with 12 CV wash buffer, elution was carried out in 6 CV elution buffer. The proteins were concentrated using Vivaspin 10k-MWCO spin concentrators (Sartorius AG, cat. no. VS15T02) up to 4 mg/mL (PdhR) and 7 mg/mL (MBP-PdhR-FUS_N_). Finally, 10% (v/v) glycerol was added and the proteins were shock-frozen in single-use aliquots in liquid nitrogen and stored at - 80 °C.

#### Formation of PdhR-FUS_N_ condensates

MBP-PdhR-FUS_N_ or PdhR proteins were thawed and desalted into the buffer used for phase separation experiments (PSB: 10 mM Tris, 150 mM NaCl, 30 mM MgCl_2_, pH 7.9) using Zeba spin desalting columns (Thermo Fisher Scientific, cat. no. 89882). MgCl_2_ was always added to the buffer directly before each experiment. The plasmids coding for SdBroccoli with different *pdhO* repeat configurations were first linearized with the *Xmn*I restriction enzyme, which has one cleavage site per plasmid. The linear DNA was isolated using the Qiagen Gel extraction kit (Qiagen, cat. no. 28704) without previously running a gel, by mixing the digestion mix with the kit’s resuspension buffer in the proportions indicated by the manufacturer. The remaining steps were done following the manufacturer’s instructions, and the elution was carried out in water. Afterwards, the buffer used for phase separation experiments was added from a 10x stock solution containing 1 M Na-pyruvate or 1 M Na-acetate. Further, MgCl_2_ was added at 30 mM directly before each experiment unless indicated otherwise. In the experiment in **Figure 3**, the protein and linearized plasmids (pCJL241) were mixed at 20 μM and 25 nM concentrations, respectively. 100 mM Na-pyruvate was added where indicated, and the same volume of buffer was added to samples without pyruvate. The protein, DNA and pyruvate mixes were incubated for 30 min at 37 °C to allow the DNA and proteins to bind. Afterwards, 3C protease was added to a final concentration of 0.1 mg/mL to induce condensate formation, and the samples were incubated for ∼20 h at RT. Subsequently, the samples were prepared for microscopy or *in vitro* transcription as described below. In experiments in **Figures 4 and 5**, the protein and linearized plasmids (pCJL240, pCJL241, pCJL244) were mixed at 20 μM and 25 nM concentrations, respectively. The protein and DNA mixes were incubated for 30 min at 37 °C to allow binding, prior to addition of 0.1 mg/mL 3C protease and incubation for ∼20 h at RT to induce condensate formation. Afterwards, Na-pyruvate or Na-acetate was added at the concentrations indicated in the figures, and the same volume of buffer was added to samples without pyruvate or acetate. The samples were further incubated for 6 h at RT. Finally, the samples were prepared for microscopy or *in vitro* transcription as described below.

**Figure 1.**
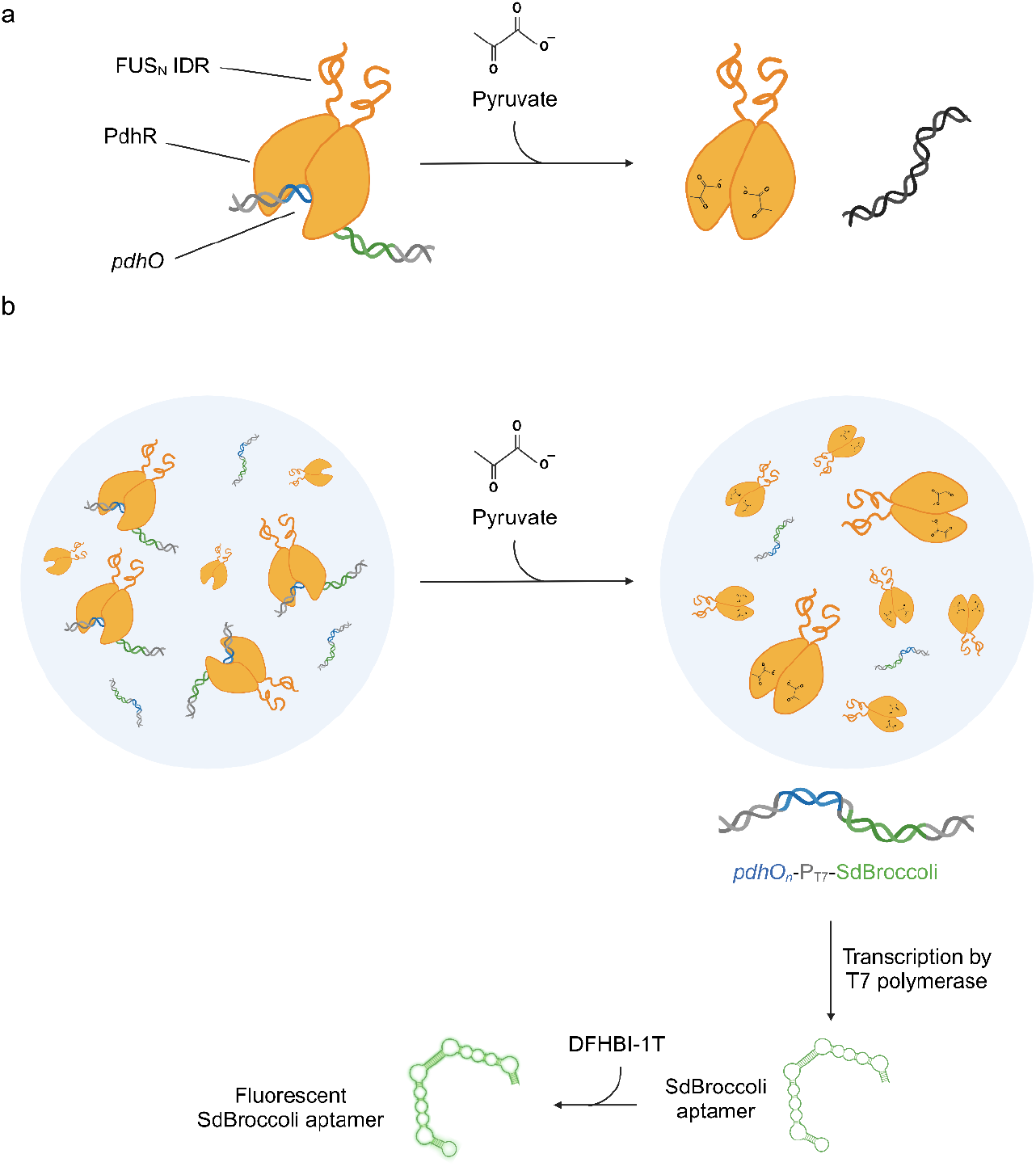
Design of the synthetic organelles with metabolite-responsive transcription control. **(a)** Molecular mechanism of metabolite responsiveness. The bacterial pyruvate dehydrogenase repressor (PdhR) binds to the pdhO sequence on a DNA molecule. Upon binding to pyruvate, the protein undergoes a conformational change that prevents it from binding to pdhO, therefore releasing the DNA molecule. **(b)** Design of the liquid-liquid phase separation-based synthetic organelles with metabolite-responsive transcriptional regulation. The pyruvate-responsive repressor protein (PdhR) is fused to the N-terminal intrinsically disordered region (IDR) of the Fused in Sarcoma protein (FUS_N_), which causes the protein to undergo liquid-liquid phase separation. PdhR recruits DNA containing the cognate *pdhO* operator which results in transcriptional inhibition of promoters contained on the construct. However, the addition of pyruvate triggers the reversible and dose-dependent release of the DNA from the coacervate which results in transcription activity from the T7 promoter. Transcription is quantified by production of the SdBroccoli RNA aptamer binding to its ligand DFHBI-1T.

**Figure 2.**
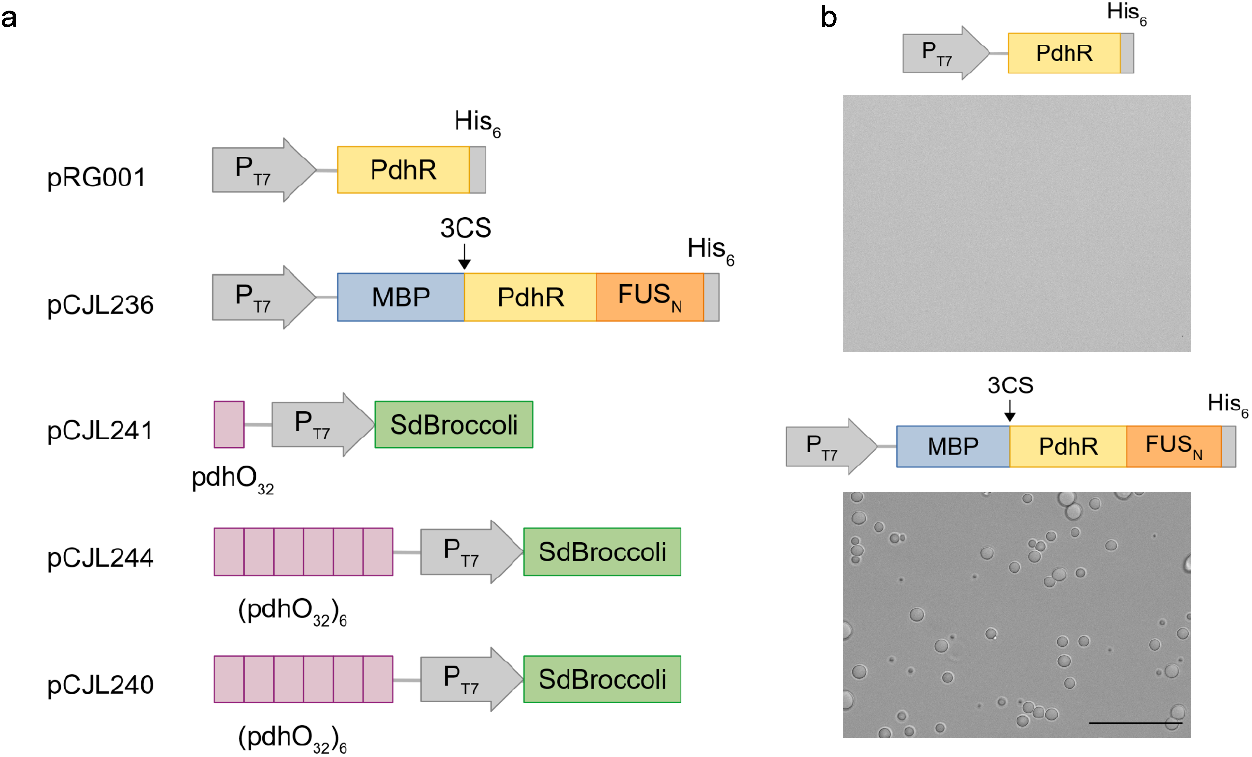
Design of the building blocks **(a)** Design of the constructs. Expression vectors for PdhR. The gene encoding PdhR was fused to the N-terminal intrinsically disordered region of FUS (FUSN_N_). To ensure solubility of the protein during production, a sequence encoding the highly soluble maltose-binding protein (MBP) was fused upstream of PdhR separated by the coding sequence for a cleavage site for 3C protease (3CS). The constructs further contained a hexahistidine-tag (His6) for purification. DNA constructs for coacervate-recruitment-responsive transcription. One (pCJL241) or six (pCJL244) repeats of the *pdhO* operator (32 bp each) were cloned upstream a polymerase T7 promoter followed by a sequence encoding the SdBroccoli aptamer. Additionally, a construct containing six truncated *pdhO* operators (17 bp each) was constructed (pCJL240). For a more detailed description of the plasmids and sequence details, see **Table S1. (b)** Coacervate formation of PdhR-FUSN_N_. 20 µM MPB-PdhR-FUSN was incubated with 3C protease to remove solubility promoting MBP prior to analysis by phase contrast microscopy (top panel). As control, a solution containing 20 µM PdhR (bottom panel) was used. Scale bar = 50 µm.

**Figure 3.**
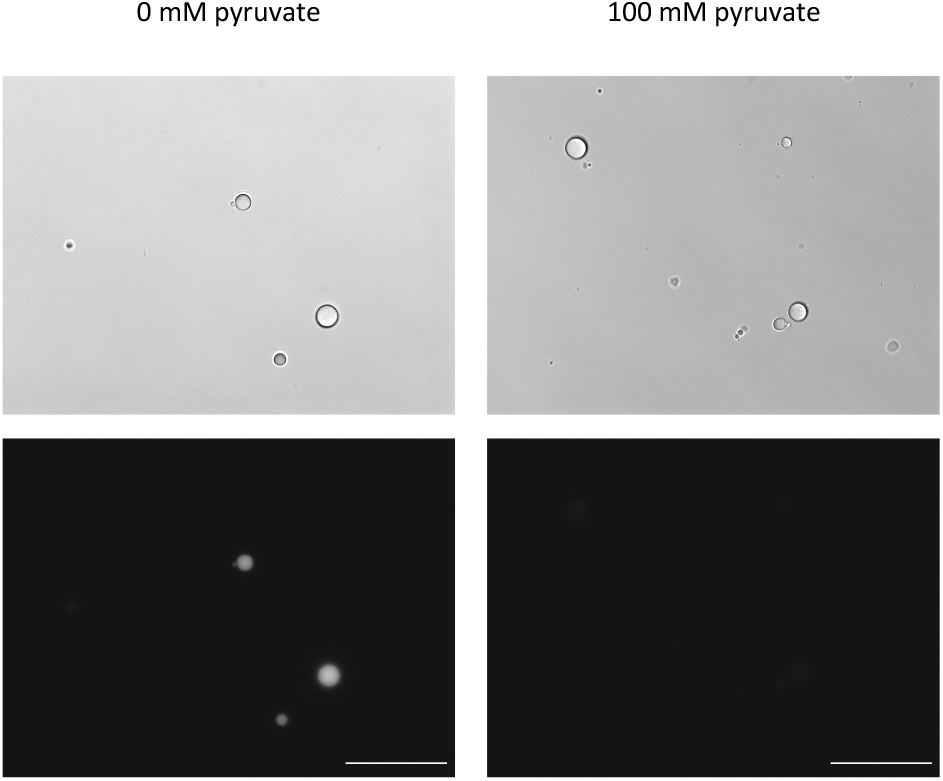
Pyruvate-responsive recruitment pdhO-containing DNA into PdhR-based coacervates. 20 µM MBP-PdhR-FUS_N_ was mixed with 25 nM linear DNA encoding *pdhO*-T_7_-SdBroccoli (pCJL241) for 30 min in the presence or absence of 100 mM pyruvate. 3C protease was added over night to remove the MBP solubility domain and to induce coacervate formation. DNA was visualized by DAPI staining and the samples were analysed by differencial interference contrast (upper panels) and fluorescence microscopy (lower panels). Scale bar = 50 μm.

**Figure 4.**
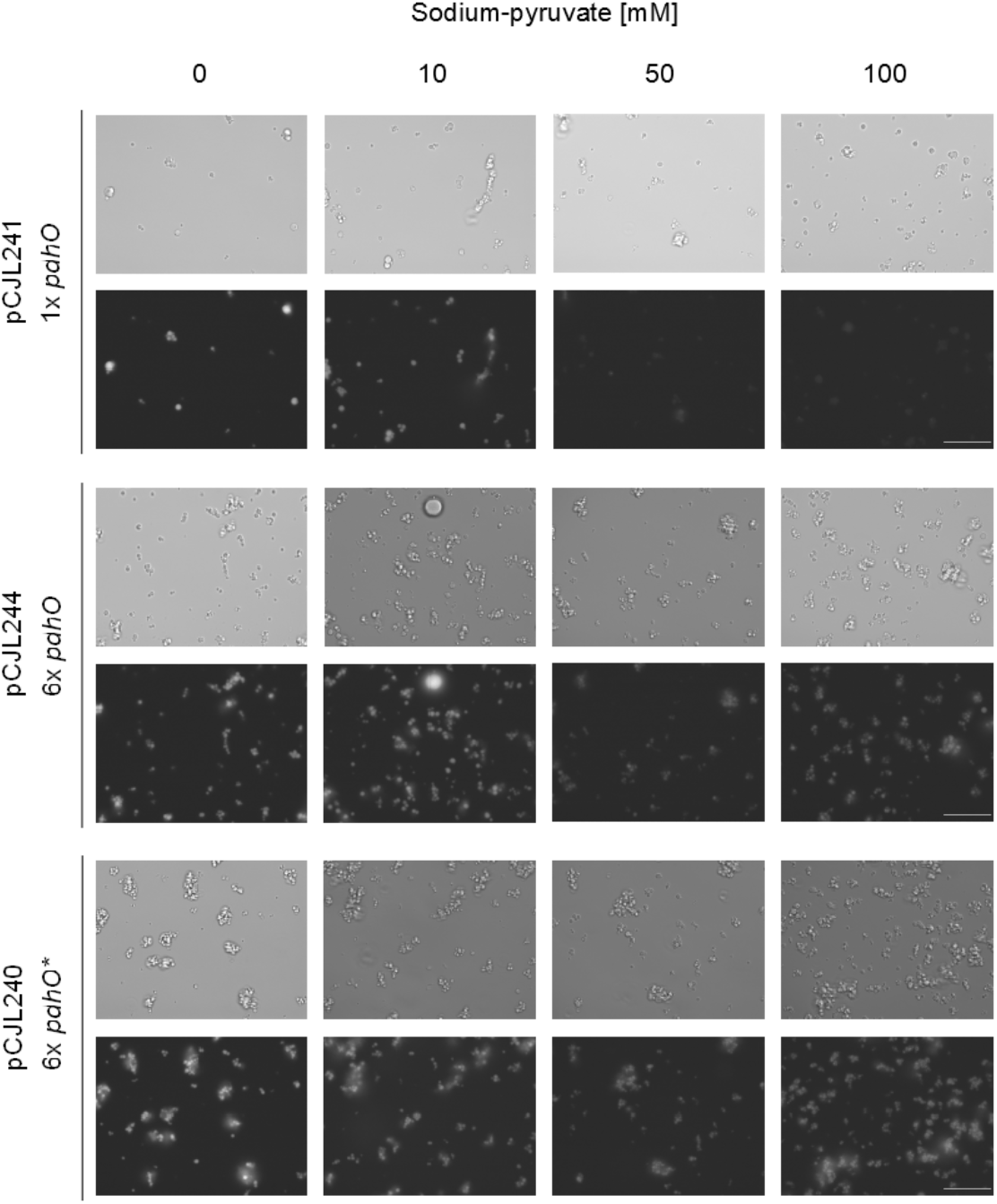
Pyruvate-reversible recruitment of *pdhO-*DNA by PdhR-FUS_N_ condensates. Linear DNA constructs with different numbers and variants of *pdhO* operators (25 nM each, see **Figure 1b**) were mixed with 20 µM MBP-PdhR-FUS_N_ prior to addition of 3C protease to induce coacervate formation by removal of the MBP tag. After incubation over night, increasing concentration of pyruvate were added. After 6 h of incubation, the samples were DAPI-stained and analysed by differencial interference contrast (upper panels) and fluorescence microscopy (lower panels). Scale bar = 50 μm.

**Figure 5.**
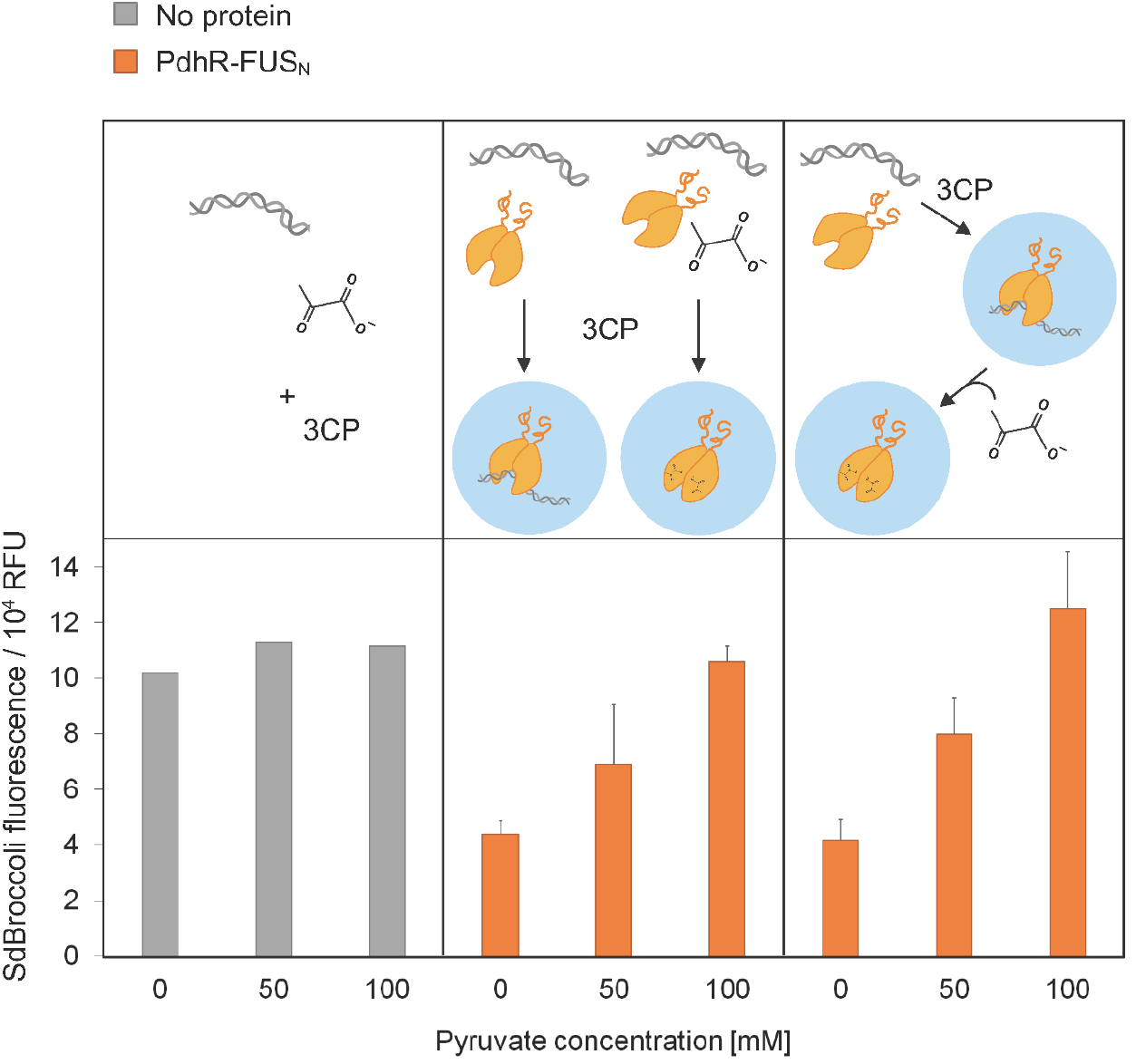
Reversible and dose-dependent control of transcription by pyruvate-responsive recruitment of DNA into PdhR coacervates. MBP-PdhR-FUS_N_ (20 µM) and a linear DNA construct (25 nM, pCJL241) encoding one *pdhO* operator upstream the T7-promoter-driven SdBroccoli expression cassette were mixed and incubated over night in the presence of 3C protease for induction of coacervate formation. Pyruvate was added either before incubation with 3CP, or added after overnight incubation at 50 mM or 100 mM concentration followed by another six hour-incubation. Subsequently, T7 polymerase and NTPs were added. After incubation over night, the SdBroccoli ligand DFHBI-1T was added and fluorescence was measured after 1 hour. As controls, samples without PhdR were included. To samples with 0 or 50 mM pyruvate, sodium acetate was added at concentrations of 100 or 50 mM, respectively, to ensure same ionic strength in all samples.

### Widefield fluorescence microscopy

To evaluate DNA sequestration into condensates, 10^−4^ mg/mL DAPI dye was added to each sample. After a 15 min incubation period, microscopy images were acquired on an AxioObserver microscope (Zeiss, Oberkochen, Germany) in differential interference contrast (DIC) mode, equipped with a Colibri 2 light source (Zeiss), a C-Apochromat 40x/1.20 W Korr objective (Zeiss) and a DAPI filter (excitation: BP 365, emission: BP 447/60-25).

### In vitro transcription

For *in vitro* transcription, a mastermix was first prepared containing transcription reagents: T7 polymerase (27 U per 100 μL reaction; Promega, Madison, WI, cat. no. P2077), 1 mM DTT and 0.5 mM rNTPs (Promega, cat. no. E6000). The reactions were incubated overnight at RT. To measure SdBroccoli RNA aptamer production, 1 mM DFHBI-1T (Bio-Techne, Minneapolis, MN, cat. no. 5610/10) was added and the samples were further incubated for 1 h at RT. Finally, SdBroccoli fluorescence was measured on a SpectraMax iD5 plate reader (Molecular Devices, San Jose, CA) using an excitation wavelength of 482 nm and detecting emission at 535 nm.

### Software

Acquisition of fluorescence microscopy image was done using the ZENblue 3.3 (Zeiss) software. All images were analyzed using FiJi ^34^. SdBroccoli fluorescent data was analyzed using Microsoft Excel 2019 (Microsoft Corporation, Redmond, WA). Figures 1 and 5b were created using BioRender.com.

## Results and discussion

For engineering a synthetic organelle with metabolite-responsive transcription control, we combined protein-based liquid-liquid phase separation with the metabolite-responsive recruitment of a transcription cassette into the formed coacervate. We hypothesized that DNA recruitment would interfere with transcription which could be reversed by metabolite-triggered DNA release. The synthetic organelle was formed by inducing LLPS of the pyruvate-responsive repressor protein PdhR^35^ via fusion to the N-terminal intrinsically disordered protein domain of FUS^36^. PdhR was shown to bind DNA constructs harbouring its cognate *pdhO* DNA operator. However, in the presence of increasing pyruvate concentrations, the DNA is released in a dose-dependent manner ^35^. As reporter for transcription from the RNA polymerase T_7_ promoter on the DNA construct, we added a coding sequence for the green fluorescent RNA aptamer, stabilized dimeric Broccoli (SdBroccoli) ^37^. Transcription was initiated by addition of T_7_ polymerase and NTPs and quantified after supplementation of the SdBroccoli ligand DFHBI-1T which emits a fluorescence signal upon complexation by the aptamer (**Figure 1**).

### Construction of the system components

First, we designed a PdhR fusion protein so that phase separation of the protein could be induced. For that purpose, the FUS IDR was fused at the C terminus of PdhR so as not to interfere with the N-terminal DNA-binding domain. To facilitate production of PdhR-FUS_N_ in soluble form, we added the Maltose-Binding Protein (MBP) solubility tag to the N-terminus separated by a 3C protease (3CP) cleavage site for later MPB removal and induction of phase separation. As control, a construct encoding PdhR alone was used. Both constructs further contained a hexahistidine tag for purification (**Figure 2a**). The constructs were produced in *E. coli* and purified by immobilized metal affinity chromatography (**Figure S1a**,**b**). The FUS_N_-containing construct was subjected to 3CP treatment (**Figure S1c**) to induce phase separation. After incubation over night, phase separation was observed with round droplets in sizes ranging from 1 to 10 μm (**Figure 2b**).

Secondly, we designed DNA constructs to be recruited into PdhR-FUS_N_ condensates. For that purpose, one or six repeats of the 32bp *pdhO* operator were placed upstream of the T7 promoter and the SdBroccoli coding sequence. We further built one construct which contained six repeats of a truncated, 17bp *pdhO* operator (**Figure 2a** and **Table S1**).

### Pyruvate-responsive recruitment of pdhO-containing DNA into PdhR-based coacervates

We analyzed the recruitment of pdhO-containing DNA into PdhR-FUS_N_ condensates in the presence or absence of pyruvate. For that purpose, MBP-PdhR-FUS_N_ was mixed with the construct containing one *pdhO* repeat (pCJL241) in the presence or absence of 100 mM Na-pyruvate. After incubation to allow for DNA and protein binding, 3CP was added to cleave the MBP tag and induce condensate formation. After incubation over night, the samples were stained with DAPI to visualize DNA localization. The samples were analysed under a fluorescence microscope with DIC contrast. As can be seen in **Figure 3**, in the absence of pyruvate, the DAPI signal colocalizes with the condensates, whereas in the presence of 100 mM pyruvate no colocalization was observed. To rule out that this effect was unspecific, e.g. due to the higher salt concentration of the pyruvate-containing sample, a control was performed with sodium acetate at the same concentration. Under these conditions, colocalization was also observed (**Figure S2**), showing that the binding and release of the DNA was specific to pyruvate.

### Reversibility and dose-dependency of pdhO DNA recruitment into PdhR-based coacervates

Having demonstrated metabolite-responsive transcription control by protein coacervates, we next evaluated the dose-dependency as well as reversibility of this effect. To this aim, we incubated the DNA constructs with different *pdhO* configurations with MBP-PdhR-FUS_N_ and removed the MBP tag by 3CP treatment to induce phase separation. After incubation over night, pyruvate was added at increasing concentrations. After incubation for 6 h, DAPI was added and the samples were analyzed by fluorescence microscope with DIC. For construct with 6 *pdhO* repeats, only partial release of the DNA was observed whereas the configuration with one single *pdhO* operator resulted in strong release at pyruvate concentrations of 50 and 100 mM thus confirming reversibility of the recruitment (**Figure 4**).

### Reversible and dose-dependent transcriptional modulation by pyruvate-responsive recruitment into coacervates

To investigate whether the pyruvate-responsive reversible DNA recruitment also correlated with reversible transcription activity from the recruited DNA, we formed coacervates containing the DNA construct with one *pdhO* operator and added increasing concentrations of pyruvate similar to the experiment above. After addition of T7 RNA polymerase and NTPs, transcription was performed over night prior to the addition of the SdBroccoli ligand DFHBI-1T and fluorescence measurement. We observed a strong drop in transcription activity in the presence of DNA-recruiting coacervates in comparison to free DNA (**Figure 5**) but also in comparison to a buffer containing non-coacervate forming PdhR at the same molar concentration as PdhR-FUS_N_ (**Figure S3**). This confirms that recruitment into the condensate rather than the binding of PdhR alone causes transcriptional downregulation. In the presence of increasing pyruvate concentrations, however, a step-wise increase in SdBroccoli fluorescence was observed indicating that the reversible release of DNA from the coacervates correlated with a dose-dependent recovery of transcriptional activity. Of note, we observed the same effect when pyruvate was added before and after coacervate formation (**Figure 5**). This data confirms the hypothesis that recruitment of the DNA construct into the coacervate-based synthetic organelle correlates with downregulation of transcription and that transcription can be reconstituted by the metabolite-responsive release in a reversible and dose-dependent manner.

## Conclusion and outlook

In this study, we have developed synthetic organelles, based on phase separation of a DNA-binding protein fused with an intrinsically disordered region, that are able of recruiting or releasing a DNA molecule in response to the absence or presence of a metabolite, and to control one of the key functions in living systems – transcription of a gene. Even though transcription-regulating phase-separated synthetic organelles have already been achieved by others ^29–31^, this is the first example in which the activity of the synthetic organelles can be regulated by a metabolite. This opens up new venues of research, such as combining the transcription-regulating synthetic organelles with other phase separation-based organelles, such as those with enzymatic activity ^23–27^, which have already been described. Ultimately, this could lead to the engineering of a fully synthetic protocell with various organelles, each performing a different function. Further, the pyruvate-responsive synthetic organelle described here can be seen as blueprint for the design of synthetic organelles responsive to a broad palette of input molecules such as different metabolites, drugs, or heavy metals by replacing PdhR with a repressor protein responsive to one of these compound classes^38,39^. The synthetic organelle approach described here may not only be instrumental for bottom-up synthetic biology in conferring minimal cells with sensing and actuation capacity but may also open new avenues in the construction of modular sensor materials for specifically detecting biomedical and environmentally important analytes.

## Acknowledgements

This work was supported by the German Research Foundation (Deutsche Forschungsgemeinschaft, DFG) under Germany’s Excellence Strategy – CIBSS, EXC-2189, Project ID: 390939984, and under the Excellence Initiative of the German Federal and State Governments – BIOSS, EXC-294 and SGBM, GSC-4.

