## Supplementary-Information for "Metabolite-responsive Control of Transcription by Phase Separation-based Synthetic Organelles"

**Table S1. Plasmids generated in this study**

| Name | Description | Sequence 5’ -> 3’ |
|---|---|---|
| **pCJL236** | $T_7$ promoter-MBP-3CS-PdhR-$FUS_N$<br>Backbone: pRG001 [33] | …GCCCTTCACCTAATACGACTCACTATAGGGGAATTGTGAGCGGATAACAA<br>TTCCCCTCTAGAAATAATTTTGTTTAACTTTAAGAAGGAATTCAGGAGCCCTT<br>CACCATGAAAATCGAAGAAGGTAAACTGGTAATCTGGATTAACGGCGATAA<br>AGGCTATAACGGTCTCGCTGAAGTCGGTAAGAAATTCGAGAAAGATACCGG<br>AATTAAAGTCACCGTTGAGCATCCGGATAAACTGGAAGAGAAATTCCCACA<br>GGTTGCGGCAACTGGCGATGGCCCTGACATTATCTTCTGGGCACACGACCG<br>CTTTGGTGGCTACGCTCAATCTGGCCTGTTGGCTGAAATCACCCCGGACAAA<br>GCGTTCCAGGACAAGCTGTATCCGTTTACCTGGGATGCCGTACGTTACAACG<br>GCAAGCTGATTGCTTACCCGATCGCTGTTGAAGCGTTATCGCTGATTTATAA<br>CAAAGATCTGCTGCCGAACCCGCCAAAAACCTGGGAAGAGATCCCGGCGCT<br>GGATAAAGAACTGAAAGCGAAAGGTAAGAGCGCGCTGATGTTCAACCTGC<br>AAGAACCGTACTTCACCTGGCCGCTGATTGCTGCTGACGGGGGTTATGCGTT<br>CAAGTATGAAAACGGCAAGTACGACATTAAAGACGTGGGCGTGGATAACG<br>CTGGCGCGAAAGCGGGTCTGACCTTCCTGGTTGACCTGATTAAAAACAAAC<br>ACATGAATGCAGACACCGATTACTCCATCGCAGAAGCTGCCTTTAATAAAGG<br>CGAAACAGCGATGACCATCAACGGCCCGTGGGCATGGTCCAACATCGACAC<br>CAGCAAAGTGAATTATGGTGTAACGGTACTGCCGACCTTCAAGGGTCAACC<br>ATCCAAACCGTTCGTTGGCGTGCTGAGCGCAGGTATTAACGCCGCCAGTCC<br>GAACAAAGAGCTGGCAAAAGAGTTCCTCGAAAACTATCTGCTGACTGATGA<br>AGGTCTGGAAGCGGTTAATAAAGACAAACCGCTGGGTGCCGTAGCGCTGA<br>AGTCTTACGAGGAAGAGTTGGCGAAAGATCCACGTATTGCCGCCACTATGG<br>AAAACGCCCAGAAAGGTGAAATCATGCCGAACATCCCGCAGATGTCCGCTT<br>TCTGGTATGCCGTGCGTACTGCGGTGATCAACGCCGCCAGCGGTCGTCAGA<br>CTGTCGATGAAGCCCTGAAAGACGCGCAGACTAATTCGAGCTCGAACAACA<br>ACAACAATAACAATAACAACAACCTCGGGATCGAGGGAAGGGGTGGAGGC<br>GGATCGCTGGAAGTTCTGTTCCAGGGGCCCATGGCCTACAGCAAAATCCGC<br>CAACCAAAACTCTCCGATGTGATTGAGCAGCAACTGGAGTTTTTGATCCTCG<br>AAGGCACTCTCCGCCCGGGCGAAAAACTCCCACCGGAACGCGAACTGGCAA<br>AACAGTTTGACGTCTCCCGTCCCTCCTTGCGTGAGGCGATTCAACGTCTCGA<br>AGCGAAGGGCTTGTTGCTTCGTCGCCAGGGTGGCGGCACTTTTGTCCAGAG<br>CAGCCTATGGCAAAGCTTCAGCGATCCGCTGGTGGAGCTGCTCTCCGACCAT<br>CCTGAGTCACAGTATGACTTGCTCGAAACACGACACGCCCTGGAAGGTATC<br>GCCGCTTATTACGCCGCGCTGCGTAGTACCGATGAAGACAAGGAACGCATC<br>CGTGAACTCCACCACGCCATAGAGCTGGCGCAGCAGTCTGGCGATCTGGAC<br>GCGGAATCAAACGCCGTACTCCAGTATCAGATTGCCGTCACCGAAGCGGCC<br>CACAATGTGGTTCTGCTTCATCTGCTAAGGTGTATGGAGCCGATGTTGGCCC<br>AGAATGTCCGCCAGAACTTCGAATTGCTCTATTCGCGTCGCGAGATGCTGCC<br>GCTGGTGAGTAGTCACCGCACCCGCATATTTGAAGCGATTATGGCCGGTAA<br>GCCGGAAGAAGCGCGCGAAGCATCGCATCGCCATCTGGCCTTTATCGAAGA<br>AATTTTGCTCGACAGAAGTCGTGAAGAGAGCCGCCGTGAGCGTTCTCTGCG<br>TCGTCTGGAGCAACGAAAGAATAAGGGCGAGCTCAATTCGAAGCTTGAAG<br>GTAAGCCTATCCCTAACCCTCTCCTCGGTCTCGATTCTACGCGTACCGGTGGT<br>GGAGGAATGGCCTCAAACGATTATACCCAACAAGCAACCCAAAGCTATGGG<br>GCCTACCCCACCCAGCCCGGGCAGGGCTATTCCCAGCAGAGCAGTCAGCCC<br>TACGGACAGCAGAGTTACAGTGGTTATAGCCAGTCCACGGACACTTCAGGC<br>TATGGCCAGAGCAGCTATTCTTCTTATGGCCAGAGCCAGAACACAGGCTATG<br>GAACTCAGTCAACTCCCCAGGGATATGGCTCGACTGGCGGCTATGGCAGTA<br>GCCAGAGCTCCCAATCGTCTTACGGGCAGCAGTCCTCCTATCCTGGCTATGG<br>CCAGCAGCCAGCTCCCAGCAGCACCTCGGGAAGTTACGGTAGCAGTTCTCA |

| | | |
|---|---|---|
| | | GAGCAGCAGCTATGGGCAGCCCCAGAGTGGGAGCTACAGCCAGCAGCCTA<br>GCTATGGTGGACAGCAGCAAAGCTATGGACAGCAGCAAAGCTATAATCCCC<br>CTCAGGGCTATGGACAGCAGAACCAGTACAACAGCAGCAGTGGTGGTGGA<br>GGTGGAGGTGGAGGTGGAGGTAACTATGGCCAAGATCAATCCTCCATGAG<br>TAGTGGTGGTGGCAGTGGTGGCGGTTATGGCAATCAAGACCAGAGTGGTG<br>GAGGTGGCAGCGGTGGCTATGGACAGCAGGACCGTGGACATCATCACCAT<br>CACCATTGAGTTTGAT… |
| **pCJL241** | *pdhO*-spacer-$T_7$ promoter-SdBroccoli<br><br>Backbone: SdB [37] | …AGCGTCTAGAGACATGAAATTGGTAAGACCAATTGACTTCGGGCTAGAGA<br>GAGCACTAACCCATCAACCTGTACGGGAACATTCTATATCGTTCTCGGACGG<br>ACAGATTACTAGAGTGCCGCTTTCAGCCCCTCTGTCGTCGCCGACGTCTGTA<br>ATATGGCGGCTAGACGATCCCGCGAAATTAATACGACTCACTATAGGAGGG<br>AGACGGTCGGGTCCATCTGAGACGGTCGGGTCCAGATATTCGTATCTGTCG<br>AGTAGAGTGTGGGCTCAGATGTCGAGTAGAGTGTGGGCTCCCTCTAGCATA<br>AC… |
| **pCJL244** | *pdhO*$_6$-spacer-$T_7$ promoter-SdBroccoli<br><br>Backbone: SdB [37] | …AGCGTCTAGAGACATGAAATTGGTAAGACCAATTGACTTCGGGCTAGAGA<br>CATGAAATTGGTAAGACCAATTGACTTCGGGCTAGAGACATGAAATTGGTA<br>AGACCAATTGACTTCGGGCTAGAGAGAGCACTAACCCGCTAGAGACATGAA<br>ATTGGTAAGACCAATTGACTTCGGGCTAGAGACATGAAATTGGTAAGACCA<br>ATTGACTTCGGGCTAGAGACATGAAATTGGTAAGACCAATTGACTTCGGGC<br>TAGAGAGAGCACTAACCCATCAACCTGTACGGGAACATTCTATATCGTTCTC<br>GGACGGACAGATTACTAGAGTGCCGCTTTCAGCCCCTCTGTCGTCGCCGAC<br>GTCTGTAATATGGCGGCTAGACGATCCCGCGAAATTAATACGACTCACTATA<br>GGAGGGAGACGGTCGGGTCCATCTGAGACGGTCGGGTCCAGATATTCGTA<br>TCTGTCGAGTAGAGTGTGGGCTCAGATGTCGAGTAGAGTGTGGGCTCCCTC<br>TAGCATAAC… |
| **pCJL240** | *pdhO**$_6$-spacer-$T_7$ promoter-SdBroccoli<br><br>Backbone: SdB [37] | …ATCGTCTAGCAATTGGTCTTACCAATTTCTAGCAATTGGTCTTACCAATTTC<br>TAGCAATTGGTCTTACCAATTTCTAGCATTTCGCGGGATCGTCTAGCAATTG<br>GTCTTACCAATTTCTAGCAATTGGTCTTACCAATTTCTAGCAATTGGTCTTAC<br>CAATTTCTAGAGAGAGCACTAACCCATCAACCTGTACGGGAACATTCTATAT<br>CGTTCTCGGACGGACAGATTACTAGAGTGCCGCTTTCAGCCCCTCTGTCGTC<br>GCCGACGTCTGTAATATGGCGGCTAGACGATCCCGCGAAATTAATACGACT<br>CACTATAGGAGGGAGACGGTCGGGTCCATCTGAGACGGTCGGGTCCAGAT<br>ATTCGTATCTGTCGAGTAGAGTGTGGGCTCAGATGTCGAGTAGAGTGTGGG<br>CTCCCTCTAGCATAAC… |

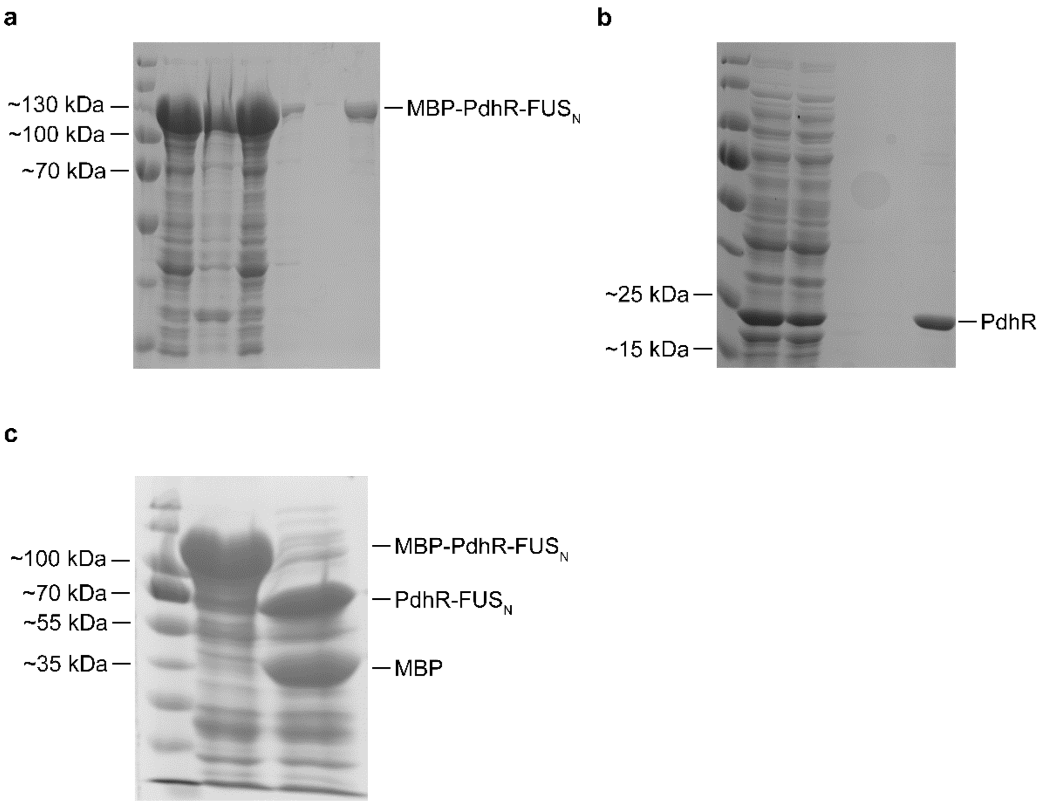


**Figure S1**. Purification and cleavage of PdhR variants. **(a)** Purification of MBP-PdhR-FUS$_N$. MBP-PdhR-FUS$_N$ encoded on plasmid pCJL236 was produced in *E. coli* and purified by immobilized metal affinity chromatography (IMAC). Samples from the different purification phases were analyzed by SDS-PAGE and Coomassie staining. Order of samples is (from left to right): protein size marker, clarified lysate, lysate pellet, flowthrough, first wash, second wash, eluate. **(b)** Purification of PdhR. PdhR encoded on plasmid pRG001 was produced and purified as described in (a). **(c)** Digestion of MBP-PdhR-FUS$_N$ with 3C protease. MBP-PdhR-FUS$_N$ (70 µM) was incubated with 2.7 µM 3C protease and incubated for 24 h at RT. Samples before (middle lane) and after (right lane) digestion were analysed by SDS-PAGE and Coomassie staining. Right lane, protein size marker.

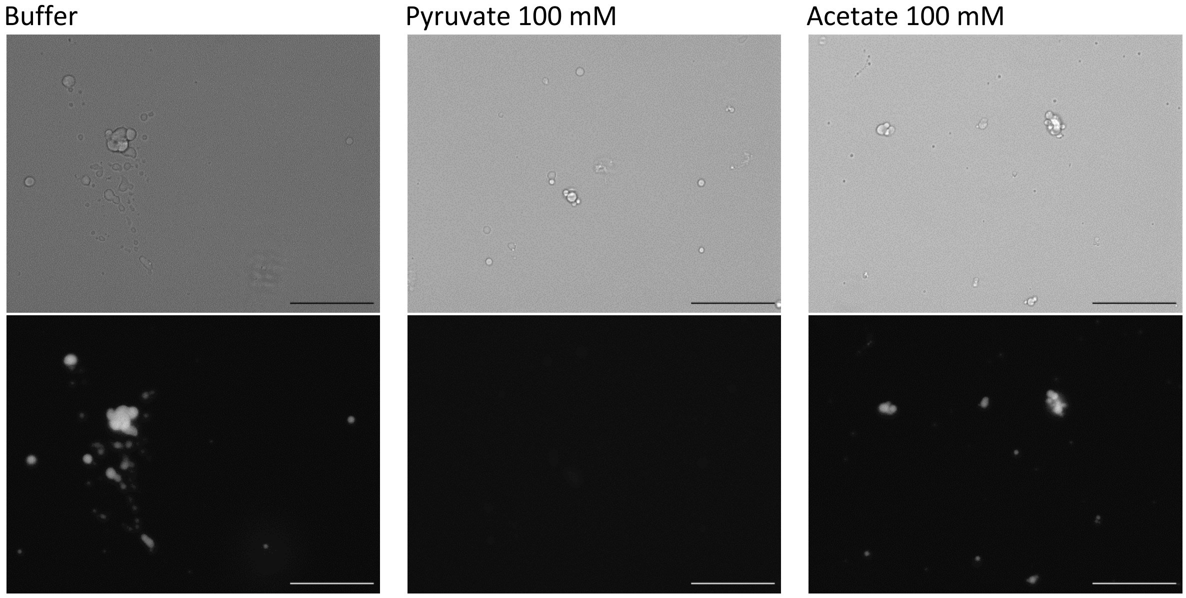


**Figure S2.** Corroborating the specificity of the effect of pyruvate on PdhR-$FUS_N$ and DNA condensates. The MBP-PdhR-$FUS_N$ protein and the pCJL241 DNA molecule were allowed to bind, and then incubated over night with 3C protease to cleave the MBP tag and induce condensate formation. Subsequently, 100 mM sodium-pyruvate or sodium-acetate were added, and the condensates were incubated for another 6 h. Finally, the samples were stained with DAPI and observed under the fluorescence microscope. Scale bar = 50 µm.

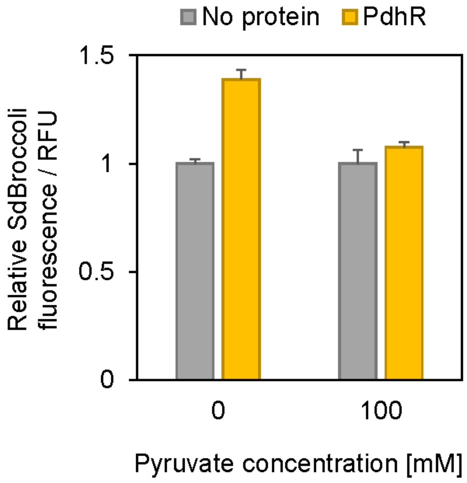


**Figure S3.** Effect of PdhR binding on SdBroccoli transcription in the presence or absence of 100 mM pyruvate. Results were normalized to the signal of the samples without protein with the same sodium pyruvate concentration.
